# A limited number of double-strand DNA breaks are sufficient to delay cell cycle progression

**DOI:** 10.1101/316158

**Authors:** Jeroen van den Berg, Anna G. Manjón, Karoline Kielbassa, Femke M. Feringa, Raimundo Freire, René H. Medema

## Abstract

DNA damaging agents cause a variety of lesions, of which DNA double-strand breaks (DSBs) are the most genotoxic. Unbiased approaches aimed at investigating the relationship between the number of DSBs and outcome of the DNA damage response have been challenging due to the random nature in which damage is induced by classical DNA damaging agents. Here we describe a CRISPR/Cas9-based system that permits us to efficiently introduce DSBs at defined sites in the genome. Using this system, we show that a guide RNA targeting only a single site in the human genome can trigger a checkpoint response that is potent enough to delay cell cycle progression. Abrogation of this checkpoint leads to DNA breaks in mitosis which give rise to micronucleated daughter cells.

## Introduction

As much as 10,000 DNA lesions arise in a human cell per day, most of which are caused by oxidative damage [1,2]. Proper management and repair of these DNA lesions is essential for development and tissue homeostasis and helps avert tumorigenesis [3–6]. Most crucial to cell viability are the pathways involved in double-strand breaks (DSBs) responses, as these represent the most genotoxic lesions [1,7]. Historically, studies aimed at a better understanding of DNA damage control have centered on the discovery of genes involved in sensitivity to DNA damaging agents [1,8]. These studies have led to the identification of a variety of damage repair pathways that act to detect and repair DNA damage. It is currently still largely unclear how these pathways act together in different genomic locations and how they are influenced by chromatin context [9,10].

Recent observations have sparked an interest in the influence of distinct chromatin states on the execution of DNA damage responses (reviewed in Sulli et al., 2012). Classic experimental approaches such as the use of DNA damaging agents like Topoisomerase II poisons or *γ*-irradiation induce breaks at random locations in the genome, making them unsuitable as tools to study site specific DSBs. Initial evidence supporting the hypothesis that local chromatin state can influence DNA damage responses has therefore come from studies using selective endonucleases, which are able to generate DSBs at single or multiple sites [12–15]. Although selective endonucleases have given us some insights regarding location-dependent effects on DNA damage responses, their applicability for unbiased investigations are limited due to a minimal regiment of target-sites in the genome (i.e. I-PpoI) or the requirement to introduce a de novo restriction site in the genome (i.e. I-SceI). Current advances in genome engineering allow us for the first time to target many, if not all, loci without the need for the introduction of de-novo sequences in the genome [16]. The genome editing technique that is currently most used is Type II clustered regularly interspaced short palindromic repeats (CRISPR), originating from a bacterial adaptive immune system that introduces DSBs in the genome of bacteriophages, thereby perturbing their bacterial virulence [17,18].

Previous work from our lab and others has shown that CRISPR can be used to tease apart location-dependent effects on checkpoints and cell fate decisions, but the systems that were used for these studies lacked sufficient temporal control over break formation [19–21]. Here, we report the generation of a time-controlled Cas9 system that allows us to induce a defined number of DSBs at very specific sites in the genome and subsequently monitor repair and cell fate. This system allows us to address how number and location of breaks influence the overall DNA damage response (DDR) and checkpoint activation.

Here we show, by using a tractable Cas9 system, that a limited number of DSBs is sensed by the DNA damage checkpoint and delay cell cycle progression.

## Results

### iCut is a tunable Cas9 system

Genome editing tools allow one to introduce DNA breaks at well-defined sites. In theory, this makes it possible to introduce a defined number of breaks in a single genome. But to render such a nuclease-based system comparable to classical DNA damaging agents it is essential to obtain temporal control over nuclease activity to prevent a cycle of cut and repair. To this end, we brought Cas9 under control of a 3rd generation doxycycline-inducible promoter [22] and introduced this into RPE-1 cells, a human diploid retinal pigment epithelial cell line immortalized by ectopic expression of hTERT [23]. While doxycycline could induce high levels of expression of Cas9 (DiC-RPE-1), we also observed considerable expression of Cas9 in the absence of doxycycline (Fig. 1B), making the system not suitable to use for time-controlled break formation. Therefore, we decided to add a destabilization domain (FKBP-degron) to the N-terminus of Cas9 and put this construct under the control of the same doxycycline-inducible promoter(Fig. 1A)[24]. Importantly, the doxycycline-regulated FKBP-tagged Cas9 (iCut) was not expressed at detectable levels in the iCut-containing RPE-1 (iCut-RPE-1) cells treated with DMSO only (Fig.1B). The addition of doxycycline (D) to the iCut-RPE-1 cells did not lead to low levels of Cas9 expression, while the combination of doxycycline and SHIELD-1 (DS) induced high levels of Cas9 expression (Fig.1B-D). Following rigorous washout, the levels of Cas9 in iCut-RPE-1 rapidly decreased (Fig. 1D). Four hours after washout, levels of expression were still near maximal in the DiC-RPE-1 cells, while they were close to basal levels in the iCut-RPE-1 cells (Fig. 1D). Additionally, we confirmed that iCut activation alone does not result in altered cell cycle distribution or proliferative defects (Suppl.Fig.1A,B). Taken together, in iCut-RPE-1 cells, Cas9 can be switched on and off in a time-controlled manner.

**Figure 1.**
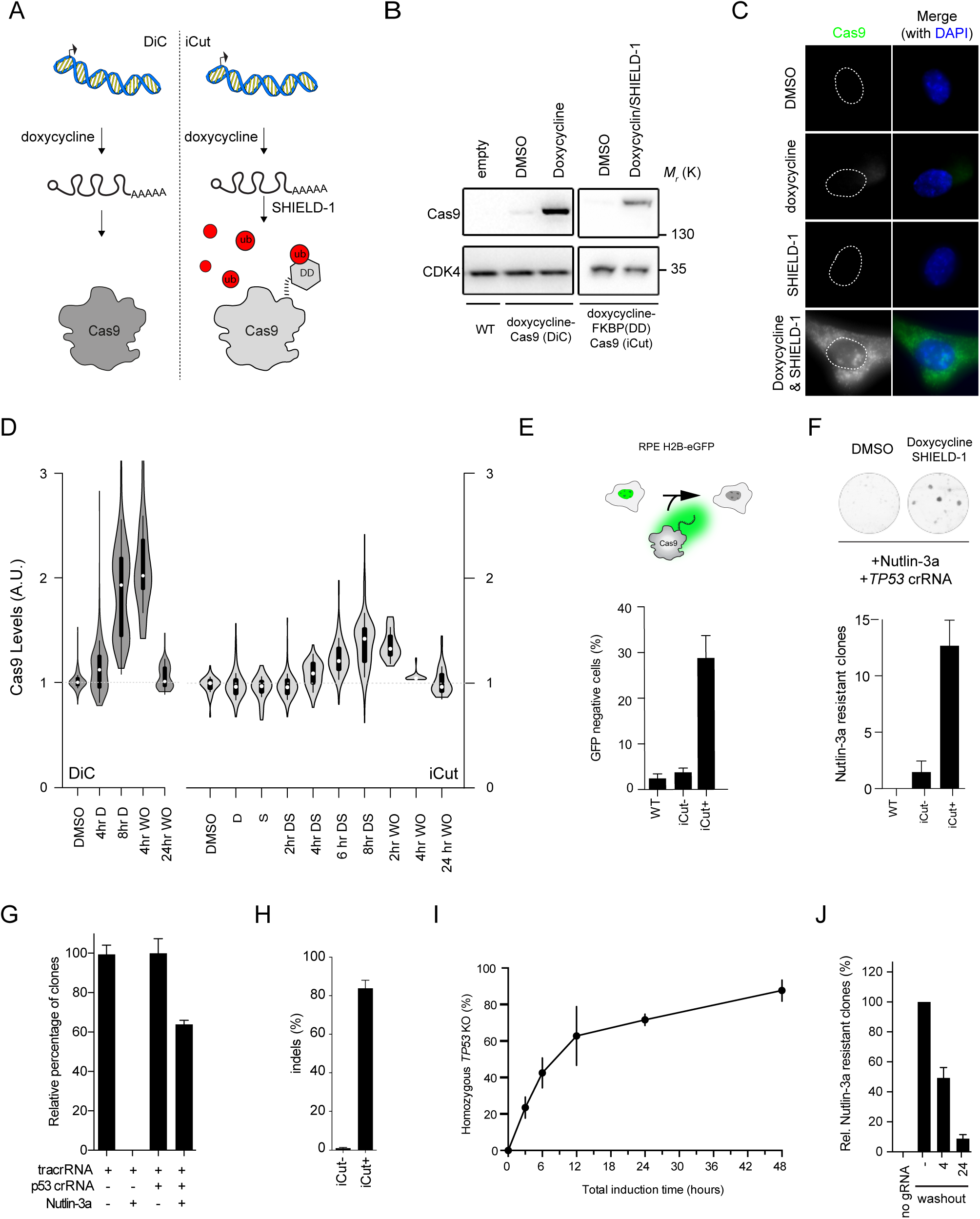
iCut is a tunable expression system for Cas9. A) Schematic representation of iCut, where levels of Cas9 can be regulated on both transcriptional and protein level. B) Western blot (WB) analysis of Cas9 expression level iCut cells after the addition ofdoxycycline and SHIELD-1 C) Immunofluorescence (IF) of iCut cells treated with DMSO (-), doxycycline (D), SHIELD-1 (S) or both agonists (DS). D) IF Quantification of Cas9 levels in DiC and iCut cells with indicated conditions. Washouts (WO) were performed after 8hrs of induction with the respective agonists. Each condition contained a least 50 cells derived from three independent. E) Top-Schematic representation of H2B-eGFP assay. Bottom - Targeting of H2B-eGFP RPE-1 cells with iCut, DS was added for 8hours following rigorous washout. Cells were allowed to recover for 4 days. eGFP+ was determined by flow cytometry (Supplemental 2B, C). The average was determined from three independent experiments (error bars represent SEM) F) Top - Clonogenic assay for *TP53* edited iCut cells. Agonists were added for 8hours following rigorous washout, cells were allowed to recover for 24 hours. Subsequently, 250 cells were plated and selected with 10μM of Nutlin-3a. Bottom - Quantification of the number of Nutlin-3a selected clones to grow out in the different conditions. G) The absolute efficiency achieved with our system by comparing Nutlin-3a with DMSO in both edited and unedited iCut cells. Agonists were added for 24 hours. The average was determined from three independent experiments (error bars represent SEM) H) The number of insertions and deletions determined by TIDE of the edited and unedited cells prior to selection. The average was determined from three independent experiments (error bars represent SD) I) Time-course of iCut activity in the presence of p53 gRNA, treated similarly to Fig.2D, *TP53KO*(%) is a ratio of Nutlin-3a/DMSO clonal outgrowth. The average was determined from three independent experiments (error bars represent SD) J) Reversibility assay of iCut system. Cells were transfected at indicated times with p53 gRNA and subsequently selected with Nutlin-3a. Timepoints indicate period after washout of doxycycline and SHIELD-1. Outgrowth was normalized to constitutive activation (-washout, error bars represent SD).

### Highly efficient gene inactivation with iCut

To functionally validate the iCut system, we first introduced a single copy of H2B-eGFP [25] into the iCut-RPE-1 cells by lentiviral transduction (Fig.1E). Subsequently, we introduced a guide RNA (gRNA) targeting the transgene and transiently activated Cas9 using the appropriate agonists for 8 hours followed by rigorous washout. Subsequently, we allowed the cells to recover for four days to allow for H2B-eGFP protein turnover. In iCut cells, we observed a loss in GFP-expression in approximately 30% of the cells (Fig.1E). The percentage of GFP-negative cells was higher in DiC-RPE-1 cells (~60%), but we also found up to 40% GFP-negative cells in the non-induced DiC-RPE-1 cultures, consistent with the observation that Cas9 is expressed in these cells in the absence of doxycycline (Suppl.Fig.2A). In sharp contrast, there was no significant increase in the percentage of GFP-negative cells in the iCut-RPE-1 treated with DMSO (iCut-), indicating that Cas9 activity is absent in iCut-RPE-1 cells in the absence of agonists (Fig.1E). Next, we monitored genome editing-efficiency of the iCut system by exploiting a small molecule inhibitor for MDM2, Nutlin-3a. Nutlin-3a inhibits proliferation of p53-proficient cells but is ineffective in p53-deficient cells [26]. Thus, only cells in which both alleles of *TP53* have been inactivated will proliferate in the presence of Nutlin3a. In accordance with previous results, we find that the highest mutation efficiency is obtained in the DiC-RPE-1 cells, but again, this system proved to be very leaky (Suppl.Fig.2B). Conversely, iCut cells display no editing activity in DMSO treated condition. However, iCut activity in the presence of doxycycline and SHIELD-1 practically reaches the levels of activated DiC-RPE-1 cells (Fig.1F).

In order to quantify the absolute activity of iCut, we transfected cells with gRNA for *TP53* or control and activated iCut for 24 hours. We allowed the cells to recover for 48 hours and plated them for clonogenic assays in the presence or absence of Nutlin-3a. Approximately 60% of the cells subjected to Cas9-induced genome-editing became resistant to Nutlin-3a, indicating that both alleles of *TP53* were inactivated in this fraction of the population (Fig.1G). Application of the TIDE assay [27] was performed in parallel, which showed that as much as 80% of the alleles had been edited in the absence of Nutlin-3a selection (Fig.1H), all of them resulting in frame-shift mutations (Suppl.Fig.2C, D). We could confirm that the editing efficiency is dependent on the time iCut is activated (Fig.1I). iCut activation for 3 hour results in ~25% Nutlin-3a-resistant cells. After 6 hr about half of the cells were able to grow out in Nutlin-3a, while at 12 hours of activation approximately two-thirds of the cells had become resistant. Longer incubation with the agonists led to a slow but consistent increase in overall genome-editing (Fig.1I), indicating that we reach near-maximum genome editing capacity after twelve hours of agonist addition. As an additional quality control for the iCut system, we set out to test reversibility of genome editing capacity following washout of the agonists. We first activated the DiC or iCut system for 16 hrs and subsequently washed out the agonists. At 4 or 24 hours following the washout of agonists we transfected the p53- or H2B-GFP-targeting gRNAs and checked genome-editing capacity. As expected, washout of doxycycline from the DiC-RPE-1 cells did not result in a decrease in activity towards *TP53* (Suppl. Fig.2E). In contrast, washout of doxycycline/SHIELD-1 led to a sharp decline in genome editing over time (Fig. 1J). Taken together, we have generated a tagged-Cas9 that allows us to regulate genome editing capacity over time.

### Introducing a defined number of breaks with iCut

We next wanted to probe whether gRNAs that target multiple homologous sites can be used in the iCut system to introduce multiple DSBs. To this end, we generated gRNAs targeting multiple sequences in the genome, which according to the CRISPOR[28] prediction tool will target 1, 4, 13, 15 or 17 homologous sites (HS1, HS4, HS13, HS15 and HS17, respectively) (Fig.2A, Suppl.Fig.3). All sites were selected based on their high on- and low off-target scores (see Materials and Methods). Using these gRNAs, we set out to investigate if the iCut system can be used to introduce an increasing number of DNA breaks. Activation of Cas9 by addition of doxycycline and SHIELD-1 occurred 16 hours prior to transfection of the gRNA. Eight hours following transfection, we fixed the cells and monitored DNA damage-induced foci using canonical DNA damage markers: 53BP1 and phosphorylated H2AX (*γ*H2AX). We find that the average number of *γ*H2AX foci ranges from 4 to 21 foci, depending on the gRNA used, whereas we find on average 3 background foci in the tracr control (Fig.2B, C). A similar number of foci are detected with 53BP1 by this set of gRNAs (Fig.2B, C). Importantly, the number of breaks that we observed seem to nicely fit with the predicted number of target sites when taking into account that a subset of cells will be in G2 (4 versus 2 alleles) and the fact that not all target sites are cut within 8 hours of activation. More accurately, we expect approximately half of the target sites to be broken around 8 hours following (see Fig.1I); so the observed numbers of breaks approximate the expected number (Fig.2B, C).

**Figure 2.**
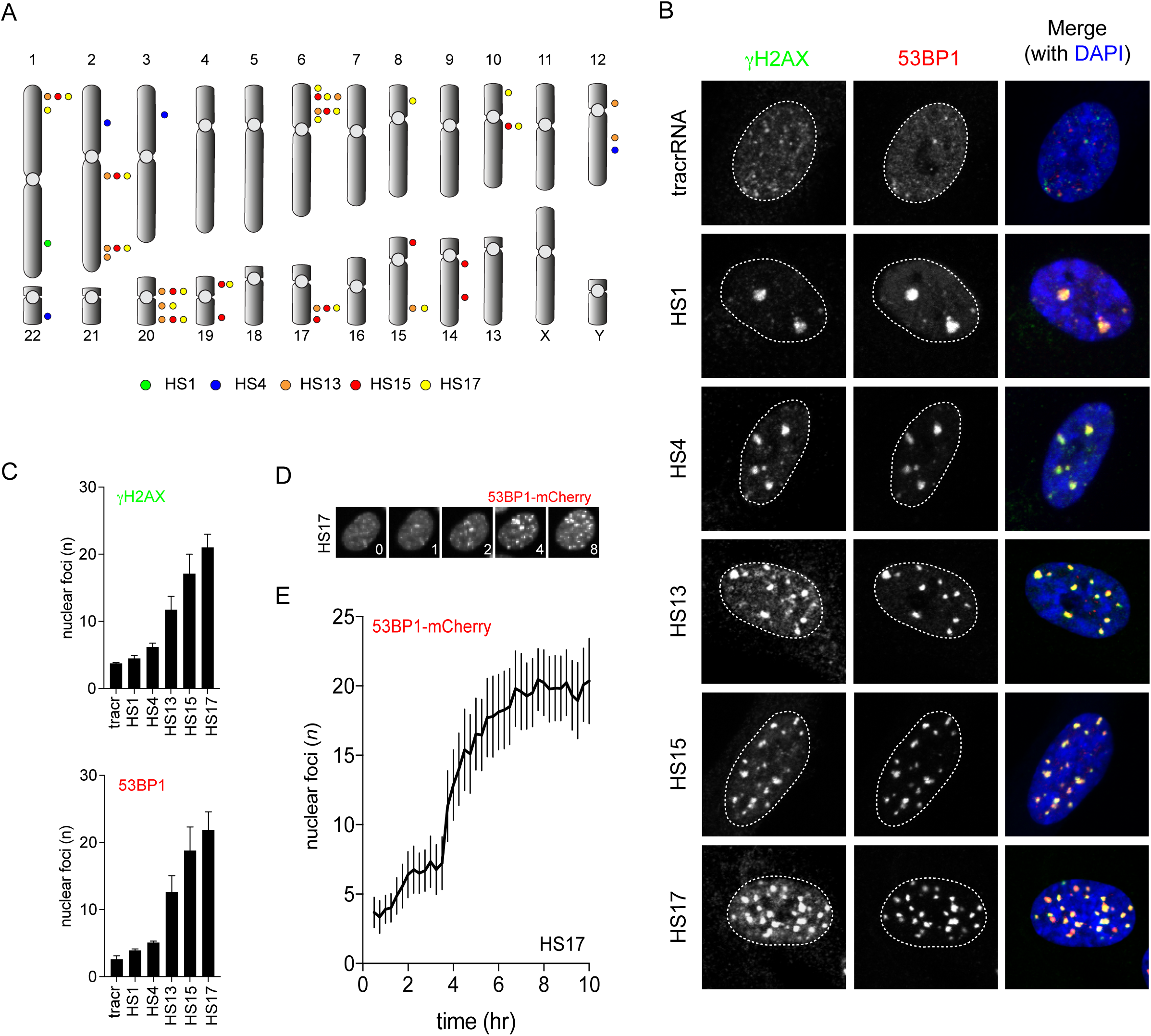
Introducing a defined number of breaks with iCut. A) Overview of genomic targeting locations with a set of gRNAs. B) iCut cells were transfected with the corresponding gRNA,16 hours later agonists were added for 8hrs. Subsequently, cells were fixed and stained with *γ*H2AX and 53BP1 to visualize DNA damage. C) Quantification of the number of nuclear foci for *γ*H2AX and 53BP1. The average was determined from three independent experiments consisting of at least 50 cells (error bars represent 95% CI). D) Representative images from 53BP1-mCherry time-lapse of iCut cells transfected with HS17 E) Quantification of the number of 53BP1-mCherry foci straight after transfecting HS17 gRNA and traced DNA damage foci formation for a time period of 10 hours. (33 cells from three independent experiment, error bars represent 95% CI)

While *γ*-Irradiation results in an instantaneous accumulation of DSBs, the timing in which Cas9 is able to target multiple loci has not been established. Therefore, we set out to approximate the time it takes for Cas9 to generate DSBs after supplying the gRNA. In order to trace the appearance of DSBs in single cells, we made use of a 53BP1-mCherry reporter. Using the HS17 gRNA in iCut cells, we observe a sharp accumulation of DSBs (Fig. 2D) at four hours post-transfection, reaching maximum break formation around 6 hours (Fig. 2E). Taken together, we are able to induce a well-defined number of DNA breaks at defined locations in single iCut RPE-1 cells with high efficiency and a relatively sharp time-resolution.

### A single break is sufficient to activate the DNA damage checkpoint

There is evidence to suggest that a single DSB is insufficient for proper checkpoint activation in mammalian cells [29,30]. On the other hand, ectopic recruitment of 100-150 identical DNA damage recognition and checkpoint proteins to a single copy LacO array is sufficient to induce a potent G2 checkpoint [31]. Additionally, a single HO nuclease-induced DSB in *Saccharomyces Cerevisiae* is able to block cells from entering mitosis, indicating that a single DSB is sufficient in yeast to induce and maintain a G2 arrest [32]. These data do not combine into a gratifying perspective on the capabilities and limitations of DNA damage checkpoints. Using the iCut system, we set out to investigate checkpoint activation at a low number of double-strand breaks. To visualize checkpoint activation, we harvest cells eight hours after transfection and investigate the levels of phospho-Ser1981 ATM and phospho-Thr68 on Chk2, a canonical ATM target[33]. The activated Chk2 kinase will, in turn, phosphorylate p53 on Ser15/20, allowing it to activate its transcriptional targets, such as the Cdk-inhibitor p21 [34–36]. Interestingly, we observe a clear induction of DNA damage signaling after the introduction of a gRNA targeting a single site (HS1) in the human genome. This is evidenced by increased phosphorylation of Ser1981 on ATM, Thr68 on Chk2 and Ser15 on p53 (Fig.3A). Thus, low levels of DNA damage (1 to 4 breaks) are able to induce ATM, Chk2 and p53 phosphorylation (Fig.3A).

**Figure 3.**
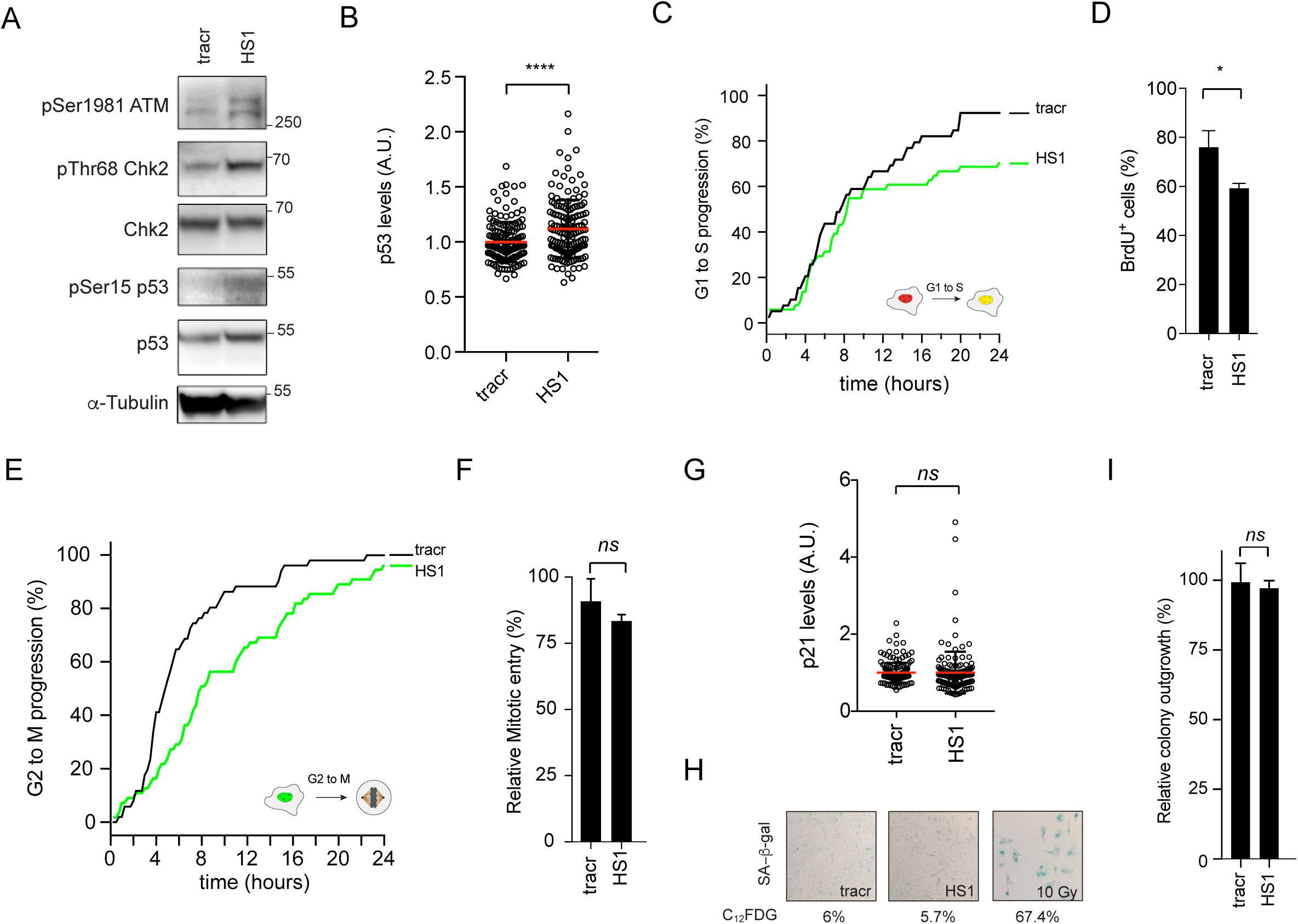
A single breaks is sufficient to activate the DNA damage checkpoint. A – Western Blot analysis of DNA damage checkpoint signaling, samples were generated in the same way as for Fig.3C B) Quantitative-IF for p53 levels 8 hours post transfected of indicated gRNAs. Student T-test was performed to asses significance. (**** = p<0.0001) C) Live cell imaging of RPE-1 FUCCI iCut cells - Cumulative S-phase entry over 24hours of RPE-1 iCut FUCCI cells challenged with indicated gRNAs. S-phase entry was quantified starting 8 hours following transfection and was determined from three independent experiments consisting of at least 50 cells. D) S-phase entry was determined by quantifying the number BrdU positive cells following sixteen our incubation. Student T-test was performed to asses significance. (**** = p<0.05) E) Live cell imaging of RPE-1 FUCCI iCut cells - Cumulative mitotic entry over 24hours of RPE-1 iCut FUCCI cells treated with indicated conditions. Mitotic entry was quantified starting 8 hours following transfection and was determined from three independent experiments consisting of at least 50 cells. F) G2 arrest following 8hrs of iCut induction in double-thymidine blocked cells with corresponding gRNA. Checkpoint recovery was assayed in the presence of Nocodazole for 16hrs cells were stained with MPM-2 to determine the mitotic percentage. The average was determined from three independent experiments (error bars represent SD) G) Quantitative-IF for p21 levels 8 hours post transfected of indicated gRNAs. Student T-test was performed to asses significance. (*ns* = not significant) H) Senescence-associated β- galactosidase staining of RPE-1 hTERT. Cells were transfected with indicated gRNAs and allowed to recover for 6 days and subsequently stained. Percentages indicate flow cytometric assessment of C_12_FDG, a fluorescent analog of SA-β-Gal. I) Clonogenic assay with 24hrs of iCut induction with the corresponding gRNA and subsequent outgrowth for 7 days. Student T-test was performed to asses significance. (*ns* = not significant).

Maintenance of a DNA damage-induced arrest relies on the induction of p53, both in G1 and in G2 [36,37], preventing cells from slipping prematurely from a checkpoint-mediated arrest [38]. We set out to investigate the upregulation of p53 in response to a gRNA targeting a single site. Using immunofluorescence, expression of p53 increases after the generation of a single break (HS1) compared to tracr control (Fig.3B). Since we observe mild but reproducible checkpoint activation with a low amount of DSBs, we next investigated how the introduction of low levels of DSBs would affect cell cycle progression. First, we interrogated the ability of cells to mount and maintain DNA damage checkpoints. Using live cell imaging of RPE-1 FUCCI cells with iCut (Fig. 3C, E); we determined the cell cycle transitions, from G1 to S and G2 to M, after introducing the HS1 gRNA. In G1 cells, we find a drop in S phase entry around 10 hr after induction of break formation. Given the fact that it takes approximately 4-6 hr for breaks to form, this suggests that the late G1 cells that have passed the restriction point fail to arrest, while the early G1 cells do arrest in response to 1 or 2 breaks (Fig.3C). Overall, the response to HS1 results in ~20% less S-phase entry over a time period of 24 hours (Fig. 3C). We were able to confirm the decrease in S-phase entry following low-level DNA damage by performing BrdU incorporation (Fig. 3D). In G2 cells, we see that mitotic entry is reduced at as little as 4 hr after break induction (Fig. 3E). Introduction of HS1 resulted in a delay in mitotic entry of about 4-6 hours, but eventually, most of the cells entered mitosis (Fig. 3E). To further corroborate these findings, DSBs were introduced in G2-synchronized cells, after which we determined the ability of cells to enter mitosis. No significant decrease in the fraction of cells entering mitosis could be observed in cells containing HS1 gRNA (Fig.3F).

Intriguingly, these data imply that cells in G1 respond differently to a single DSB when compared to cells in G2. When the damage is encountered in G1, cells readily enter S-phase up to 10 hours following DSB formation. After this period, inactivation of the G1 checkpoint is hindered as only a couple of cells are still able to enter S-phase beyond this point in time. Thus, cells that are in late G1 are incapable of mounting a checkpoint that prevents S phase entry, whereas the early G1 cells can. This late G1 population has previously been described to be beyond the restriction point and therefore unable to arrest in G1[39–41]. Conversely, the G2 checkpoint reacts much more rapidly to a single DSB, by blocking the progression into mitosis as early as 4 hr after break formation. However, in contrast to the prolonged G1 arrest, the G2 arrest is much more transient in nature. Over time, cumulative mitotic entry catches up to the unperturbed cells in the HS1-treated condition, indicating that (almost) all of the arrested cells can revert the cell cycle block in 4-6 hr (Fig. 3E). Taken together, these data show that cells with a minimal amount of DNA breaks are capable to mount an effective checkpoint response. The type of response depends on which phase of the cell cycle the damage is encountered. The deterministic factors distinguishing an arrest (G1) versus a delay (G2) remain to be uncovered, but iCut allows us to rule out location or number as a possible explanation for this difference. Also, it should be noted that the prolonged arrest in G1 is induced by 1-2 breaks, while the reversible arrest in G2 is induced by 1-4 breaks.

To investigate whether the HS1-induced cell cycle arrest can result in a permanent withdrawal from the cell cycle, we first quantified p21 expression levels. Strikingly, we observe no upregulation in cells challenged with a single DSB (Fig. 3G). This is counter-intuitive as we do observe an upregulation of p53 levels (Fig. 3B). The absence of p21 in the presence of breaks and p53 upregulation suggests a lack of cell cycle exit [42,43]. To address cell cycle exit directly, we quantified the fraction of senescent cells seven days following the initial DNA damage insult. Cells treated with HS1 did not display an increased propensity to enter senescence (SA-β-Gal, Fig.3H). We were able to validate these results by means of flow cytometric analysis of C_12_FDG, a fluorescent analog of β-Gal (Fig. 3H & Suppl. Fig. 4). Conversely, cells challenged with 10Gy of irradiation readily converted to senescent cells (SA-β-Gal) and approximately 75% of the cells became C_12_FDG positive (Fig. 3H).

Additionally, we performed clonogenic assays to monitor the ability of cells to grow out after DSB formation. Using HS1, we find a minor decrease in colony outgrowth ~5% compared to control (tracrRNA alone) cells, indicating this number of breaks is not enough to cause a significant proliferative disadvantage (Fig.3I). Irradiating cells with 10Gy of irradiation invariably led to zero colonies (data not shown). Taken together, we observe that DNA damage signaling is activated upon a single DSB. This signaling is enough to halt cells for a different period of time in the G1 versus the G2 phase of the cell cycle.

However, the induced arrest did not lead to a permanent cell cycle exit, indicating that cells are able to respond to very low levels of genotoxic stress but cope with this without removing themselves from the proliferating population.

Future studies are required to identify the contribution of a single break among multiple DSBs to checkpoint signaling and establishment. To investigate this, sensitive biosensors of checkpoint activation (such as the ATM and ATR activity probe [44]) should be employed. Work from the Lahav lab has shown that cell fate dictated by the DNA damage response is in part regulated by the manner in which p53 transcriptional activity is induced [45]. More recent work from this group showed that p53 levels need to rise above a threshold to execute apoptosis, but that this threshold increases with time after initial break formation [46]. Thus, in order to induce a reversible cell cycle arrest, it is of the utmost importance to strictly keep p53 activity below the threshold required to induce apoptosis [46] or senescence [42,47,48]. In light of results described here, it would be interesting to investigate the relationship between DNA checkpoint activation at specific locations and cell fate decisions governed by p53 signaling. Moreover, persistent breaks could lead to a longer or higher induction of damage-induced transcription leading to different outcomes which might be dependent on break location. Thus, we expect that the outcome of the response depends on more than just numbers of breaks and will also be dependent on the ease with which breaks are repaired, but more work is required to resolve this issue.

### Abrogation of a single DSB induced G2 checkpoint causes genomic instability

It has been extensively shown that cells readily enter into mitosis in the presence of a low number of breaks[29,44,47]. Therefore, our observation that cells display a checkpoint-induced delay in G2 in response to a low number of DSBs is quite unexpected and contradicts the current consensus in the field, stating that the G2 checkpoint is incapable of sensing a low number of breaks. However, our data thus far do not resolve if the delay is functional. In other words, is the delay required to allow time for DNA damage responses and faithful repair to occur? To address this, we asked if an override of this checkpoint response would comprise genomic integrity. To abrogate the activity of the G2 checkpoint we used two different settings, dual inhibition of ATM and ATR (AT2i) or inhibition of Wee1 (Wee1i). We used RPE-1 iCut FUCCI cells to trace single cells in which we assessed mitotic entry following break induction using HS1. Again, we observed a delay of approximately 4-6hrs in the HS1-transfected cells compared to tracr-transfected cells (Fig. 4A, DMSO). Conversely, when we treated cells with either AT2i or Wee1i, the delay was drastically shortened (Fig. 4A, AT2i & Wee1i). Next, we asked if this checkpoint override will cause cells to enter mitosis with residual DNA breaks (Fig. 4B, C). Indeed, upon abrogation of the G2 checkpoint, HS1-transfected cells that had entered mitosis stained positive for *γ*H2AX and MDC1 foci (Fig. 4C, D). This indicates that the G2 checkpoint is required to prevent cells from entering mitosis, even if they only have a very limited number of breaks. Next, we wondered what the consequence is for cells when we ablate the HS1-induced G2 checkpoint. To our surprise, we observed a 5-fold increase in the frequency of micronuclei generated after overriding the G2 checkpoint by either AT2i or Wee1i (Fig.4E, F). This indicates that the checkpoint response in G2, triggered by 1-4 breaks is essential to prevent cell division in the presence of a broken chromosome. Since it was previously reported that the inheritance of a broken chromosome can severely affect cell viability[49], we subsequently analyzed the proliferative capacity of G2 synchronized HS1-treated cells with or without a functional G2 checkpoint. We find that in cells with a functional G2 checkpoint, viability is not affected, as evidenced by clonogenic outgrowth (Fig.4G). Consistent with our observation that HS1-transfected cells treated with AT2i or Wee1i slip into mitosis with DNA breaks, ablation of the G2 checkpoint hampers proliferation and outgrowth in HS1-transfected cells (Fig. 4G). Taken together, a low number of breaks in G2 will activate a checkpoint response that is absolutely required to prevent the propagation of DNA lesions into mitosis and the subsequent generation of micronucleated cells, a hallmark of genomic instability.

**Figure 4.**
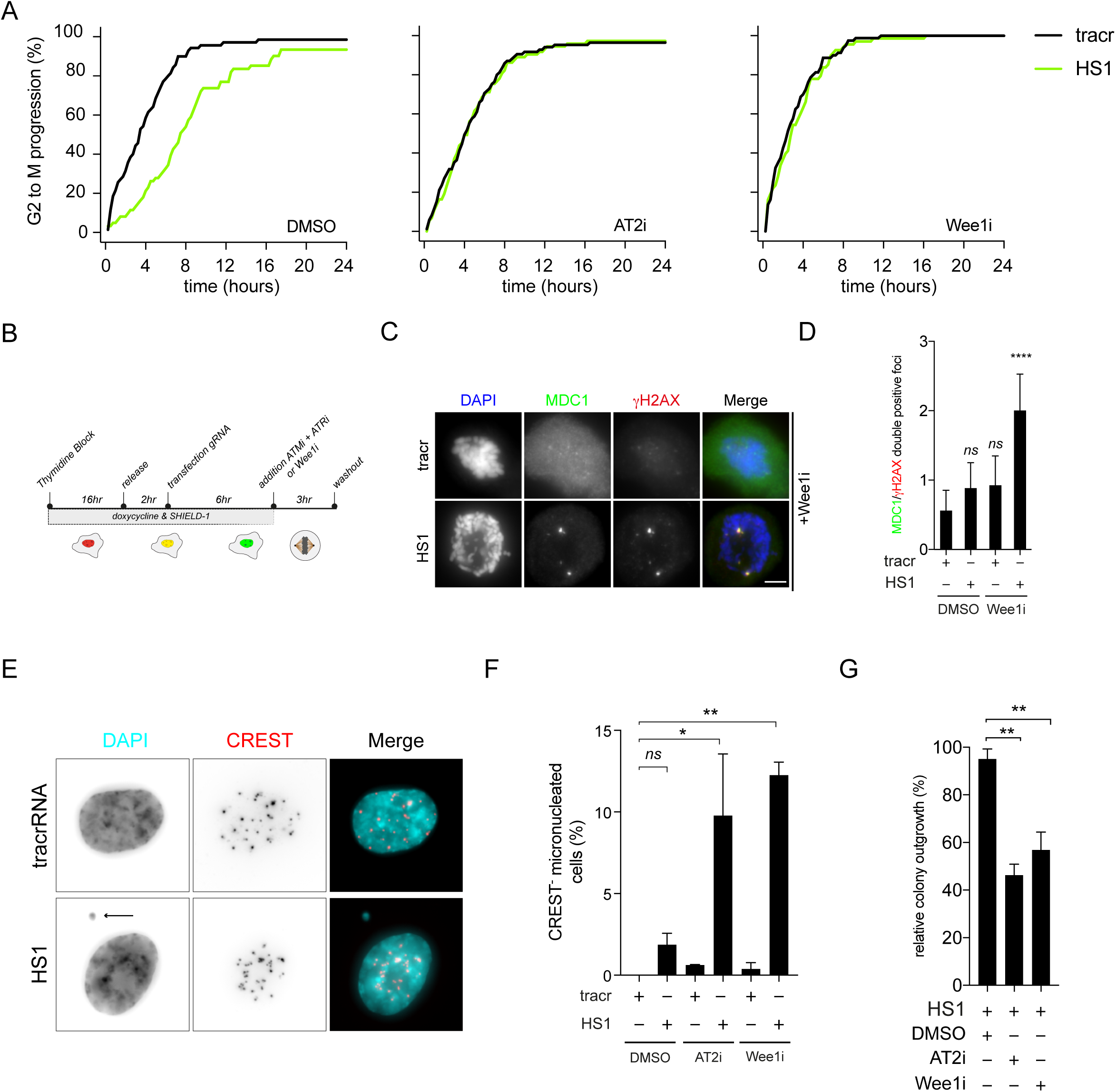
G2 checkpoint is essential to prevent genomic instability and maintain fitness in the presence of low numbers of breaks. A) RPE-1 FUCCI iCut cells transfected with HS1 or tracr in the presence or absence of either AT2i (ATMi and ATRi) or Wee1i. Mitotic entry was quantified starting 8 hours following transfection and was determined from three independent experiments consisting of at least 50 cells. B) Schematic overview of G2 checkpoint ablation in the presence of low number of DNA breaks C) Nocodazole trapped mitotic cells transfected with HS1 or tracr in the presence of a Wee1i, stained for MDC1 and *γ*H2AX. D) Quantification of Fig. 4C. One way ANOVA with Bonferroni’s multiple comparisons test was performed to asses significance. (*ns* = not significant, **** = p<0.0001) E) Nuclear morphology of cells treated with AT2i transfected with tracr or HS1. F) Quantification of the phenotype observed in Fig. 4E. One way ANOVA with Bonferroni’s multiple comparisons test was performed to asses significance. (*ns* = not significant, * = p<0.05. ** p < 0.005) G) Clonogenic outgrowth of cells transfected with HS1 normalized by tracr alone in corresponding conditions. One way ANOVA with Bonferroni’s multiple comparisons test was performed to asses significance. (*ns* = not significant, ** = p<0.005)

These findings indicate that a single DSB has the potential to mount a DNA damage checkpoint. Particularly, the G2 checkpoint in this setting displays interesting characteristics such as a delay dependent on DNA damage checkpoint kinases and a surprising impact on genomic stability when abrogated. In light of these findings, it would be intriguing to assess whether responses are regulated in another fashion once we target different regions in the genome. One could hypothesize that differences occur once DSB targeting occurs more distal or proximal with respect to the centromere. Another exciting line of investigating would be to target DSBs in a region with high or low transcriptional activity. These options are now feasible by using CRISPR/Cas9 mediated breaks.

Finally, the DNA repair and damage response fields have generated many reporter systems based on integrating an I-SceI site into the genome couple to a functional read-out[50]. These readouts vary from assaying DNA damage-induced inhibition of transcription in neighboring genes to quantifying the relative rate of either HR or NHEJ repair pathway[15,51,52]. Initially, these systems were utilized to validate novel players in these responses. We would like to propose using this CRISPR system to induce DSBs at specific locations to determine the contribution of DNA sequence and chromatin in DDR and repair phenotypes.

## Materials and Methods

### Antibody generation

Anti-Cas9 was raised against the first 300 amino acids of Cas9 from *Streptococcus pyogenes*. The cDNA corresponding to amino acids 1-300 of Cas9 from *S. Pyogenes* was cloned in pET-30a (Novagen). The resulting 6x His tagged antigen was expressed in *Escherichia Coli*, purified and used for rabbit immunization. Rabbit polyclonal antiserum was affinity purified.

### Plasmid Construction

pCW-Cas9 was a gift from Eric Lander and David Sabatini (Addgene plasmid # 50661). The iCut plasmid was generated by linearizing pCW-Cas9 with NheI and introducing the FKBP destabilization domain. The FKBP destabilization domain was amplified from the pRetroX-Tight FKBP-I-PpoI plasmid (Warmerdam et al. 2016) using primers suitable for Gibson Assembly (Gibson et al. 2009).

### Cell lines and Tissue Culture

Retinal pigment epithelial (RPE-1) hTERT cell lines (obtained from American Type Culture Collection) were maintained in DMEM/F12 GlutaMAX medium (Gibco) containing 10% Fetal Bovine Serum and 1% Penicillin-Streptomycin (10.000U/mL). DiC-RPE-1 cells were generated by lentiviral transduction with 1^st^ generation lentiviral helper plasmids of pCW-Cas9 [22]. iCut-RPE-1 cells were generated by transduction of the iCut plasmid, followed by selection with Puromycin (20μg/mL). Constitutive Cas9 expressing cells were previously described [20]. Chemicals used in this study: Doxycycline (Sigma, 1mM), SHIELD-1 (Aobious, 1μM), Nutlin-3a (Cayman Chemical, 10μM), Nocodazole (250 μM, Sigma).

### tracrRNA:crRNA design and transfections

Edit-R crRNA were designed with on-target scores determined by the Rule Set 2 [54]. Off-target scores were determined by the specificity score [55]. For HS1, we used the previously described crRNA targeting the *LBR* gene from Brinkman et al. 2014. For HS4, we used a sequence of the *NF2* gene and processed similarly as HS13 and HS18 to select a crRNA with the most target sites. For HS13, HS15 and HS17; we used *RPL12* pseudogenes to design sgRNAs with the rationale that these would target multiple sequences. We used the *RPL12P38* pseudo-gene annotated in the hg19 assembly of the human genome. Subsequently, we selected sgRNAs based on the MIT CRISPR prediction tool (crispr.mit.edu). We included predicted sites with complete homology and with maximum 1 mismatch outside of the seed sequence of the sgRNA (position 1-8,Semenova et al. 2011). Out of all the targets none target coding sequences of genes. tracrRNA:crRNA duplex was transfected according to manufacturer’s protocol with one exception [57]; we used RNAiMAX according to manufacturer’s protocol, instead of DharmaFect Duo, as we did not transfect plasmid DNA.

The following crRNA were used in this study:

~~~
eGFP 5′-GTCGCCCTCGAACTTCACCT-3′ Doench 2016 [70], Hsu 2013 [81]
*TP53* 5′- TCGACGCTAGGATCTGACTG-3′ Doench 2016 [64], Hsu 2013 [48]
HS1 5′-GCCGATGGTGAAGTGGTAAG-3′ Doench 2016 [73], Hsu 2013 [55]
HS4 5′- TGGACTGCAGTACACAATCA-3′ Doench 2016 [58], Hsu 2013 [16]
HS13 5′-AGAAAAACATTAAACACAGT -3′ Doench 2016 [58], Hsu 2013 [6]
HS15 5′-TTTTTGGAGACAGACCCAGG-3′ Doench 2016 [77], Hsu 2013 [5]
HS17 5′-CAGACAGGCCCAGATTGAGG-3′ Doench 2016 [70], Hsu 2013 [4]
~~~

### Clonogenic assays

iCut-RPE-1 or DiC-RPE-1 cells were transfected with the indicated crRNAs and 16 hours later, 250 single cells per well were seeded in 6 well plates. Cells were treated with the indicated drugs and allowed to grow out for 7 days. Plates were fixed in 80% ice-cold Methanol and stained with 0.2% Crystal Violet solution. Colonies were counted and normalized to plating efficiency of untreated control.

### Determination of insertions and deletion by TIDE

Insertions and deletion of the *TP53* locus were quantified by PCR amplification of the edited region with the following primer (5’-gggaaggttggaagtccctctc-3’ and GCTTCATCTGGACCTGGGTCTT). This PCR product was subjected to Sanger Sequencing and analyzed by the Tracking of Indels by Decomposition (TIDE) method [27].

### Fixed Cell Microscopy

Images were obtained using a DeltaVision Elite (Applied Precision) maintained at 37°C equipped with a 60× 1.45 numerical aperture (NA) lens (Olympus) and cooled CoolSnap CCD camera. DNA damage foci were evaluated in ImageJ, using an in-house developed macro that enabled automatic and objective analysis. In brief, cell nuclei were detected by thresholding on the (median-filtered) DAPI signal, after which touching nuclei were separated by a watershed operation. The foci signal was background-subtracted using a Difference of Gaussians filter. For every nucleus, foci were identified as regions of adjacent pixels satisfying the following criteria: (i) the gray value exceeds the nuclear background signal by a set number of times (typically 2-fold) the median background standard deviation of all nuclei in the image, and is higher than a user-defined absolute minimum value [1]; (ii) the area is larger than a defined area (typically 16 pixels). These parameters were optimized for every experiment by manually comparing the detected foci with the original signal.

### Live-cell imaging

Cells were grown in Lab-Tek II chambered coverglass (Thermo Scientific) in Dulbecco’s Modified Eagle Medium/ Nutrient Mixture F-12 (DMEM/F12) outfitted with a CO2 controller set at 5%. Images were obtained using a DeltaVision Elite (Applied Precision) maintained at 37°C equipped with a 10× or 20× PLAN Apo S lens (Olympus) and cooled CoolSnap CCD camera. Macro’s used to quantify foci living cells were previously described in Feringa et al. 2016.

### FACS analysis

For G2 checkpoint recovery, we synchronized cells at the G1/S transition using a thymidine block, we induced Cas9-activity by the addition of doxycycline/SHIELD-1 at the time of thymidine addition. At the time of release, the cells were transfected with indicated gRNAs and the cells were allowed to progress into G2. Nocodazole was added 7 hours post transfection to trap cells in mitosis over a period of 16 hours. Cumulative mitotic entry was determined by the percentage of MPM-2-positive cells [58]. For H2B-eGFP disruption, we fixed cells in 70% ethanol 4 days after the washout of the agonists (doxycycline for DiC, doxycycline, and SHIELD-1 for iCut). Disruption was analyzed by determining the percentage eGFP negative cells in the total population.

### Immunofluorescence and Western Blots

For IF, cells were fixed with 3.7% formaldehyde for 5 min and permeabilized with 0.2% Triton-X100 for 5 min before blocking in 3% bovine serum albumin (BSA) in PBS supplemented with 0.1% Tween (PBS-T) for 1 h. Cells were incubated overnight at 4°C with primary antibody in PBS-T with 3% BSA, washed three times with PBS-T, and incubated with secondary antibody and DAPI in PBS-T with 3% BSA for 2 h at room temperature (RT). Western Blot analysis was performed as previously described (Feringa et al. 2016). The following primary antibodies were used in this study: anti-Cas9 (homemade rabbit polyclonal, 1:2000), anti-MPM2 (05-368, Millipore, 1:500), anti-*γ*H2AX (ser139p; 05–636 Upstate, 1:500), anti-53BP1 (H-300, Santa Cruz, sc-22760, 1:500) anti-Cdk4 (C-22; sc-260 Santa Cruz, 1:1000), anti-pSer15 p53 (Cell Signaling, #9286, 1:250), anti-p53 (DO-1, Santa Cruz, sc-126, 1:500), anti-Chk2 (H-300, Santa Cruz, sc-9064, 1:500), anti-pThr68 Chk2 The following secondary antibodies were used for western blot experiments: peroxidase-conjugated goat anti-rabbit (P448 DAKO, 1:2000) and goat anti-mouse (P447 DAKO, 1:2000). Secondary antibodies used for immunofluorescence and FACS analysis were anti-rabbit Alexa 488 (A11008 Molecular probes, 1:600), anti-mouse Alexa 568 (A11004 Molecular probes, 1:600). DAPI was used at a final concentration of

### Senescence-associated beta-galactosidase (SA-β-gal) activity assays

To detect SA-β-gal by cytochemistry, cells were fixed for 5 min using 2% formaldehyde 0,2% gluteraldehyde in PBS after 6 days of gRNA transfection. Cells were washed three times with PBS before overnight (16 h) incubation in staining solution (X-gal in dimethylformamide (1 mg ml), citric acid/sodium phosphate buffer at pH6 (40 mM), potassium ferrocyanide (5 mM), potassium ferricyanide (5 mM), sodium chloride (150 mM) and magnesium chloride (2 mM)) at 37°C (not in a CO_2_ incubator). Cells were washed with PBS and the blue staining was detected using a CCD microscope equipped with a Zeiss AxioCam colour camera (Axiocam HRc). The C_12_FDG assay is based on the hydrolysis of a membrane permeable molecule, the 5-dodecanoylaminofluorescein di-β- D-galactopyranoside (C_12_FDG), by β-galactosidase enriched in senescent cells. After hydrolysis and laser excitation, the C_12_FDG emits green fluorescence and can therefore be detected by flow cytometry. After 6 days of gRNA transfection, RPE-1 were incubated with the Bafilomycin A1 solution for 1 hour at 37°C, 5% CO_2_ followed by the C_12_FDG staining for 2 hours at 37°C, 5% CO_2_. After incubation, RPE-1 were trypsinized and resuspended with PBS. Cells were analyzed immediately using FACS Calibur flow cytometry. C_12_FDG was measured on the FL1 detector.

## Acknowledgements

The Dutch Cancer Foundation (KWF; NKI 2014–6787), the Netherlands Organization for Scientific Research (NWO) (022.001.003) and Top-Go ZonMw (91210065) funded this research. We thank Bas van Steensel, Eva Brinkman, Joppe Nieuwenhuis (Netherlands Cancer Institute) and Aniek Janssen (Lawrence Berkeley National Laboratory) for advice and critical reading of the manuscript. We thank all Medema, Rowland and Jacobs lab members for the helpful discussions on this study.

## Author contributions

J.vd.B. and R.H.M. conceived and designed the experiments, and wrote the paper. J.vd.B., A.G.M., F.M.F. & K.K. performed the experiments and analyzed the data. R.F. generated the polyclonal rabbit-anti-Cas9 antibody and performed the initial validation.

## Conflict of interest

The authors declare that they have no conflict of interest

---

**Supplemental Figure 1.**
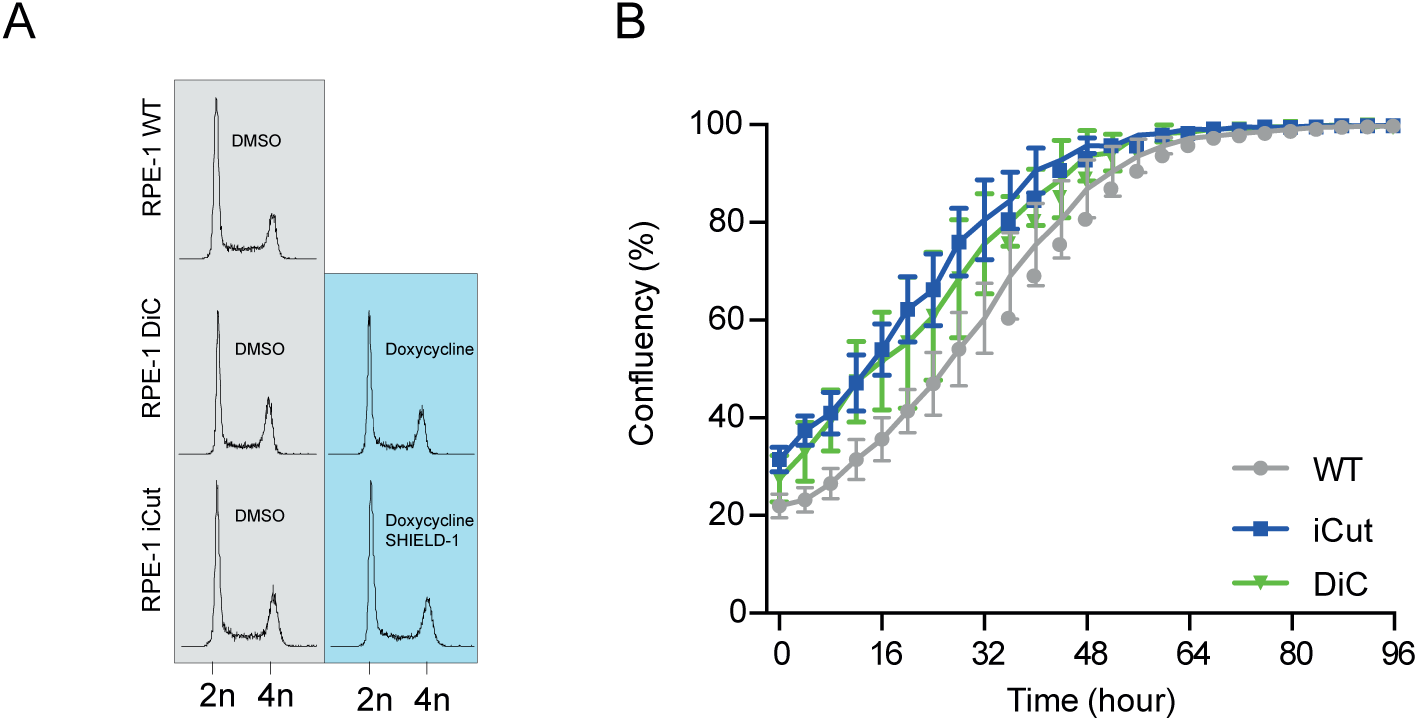
A) Cell cycle profile of RPE-1 WT, RPE-1 DiC and RPE-1 iCut analyzed by Flow Cytometry. B) Growth curve of RPE-1 WT, DiC and iCut cells in the presence of agonists.

**Supplemental Figure 2.**
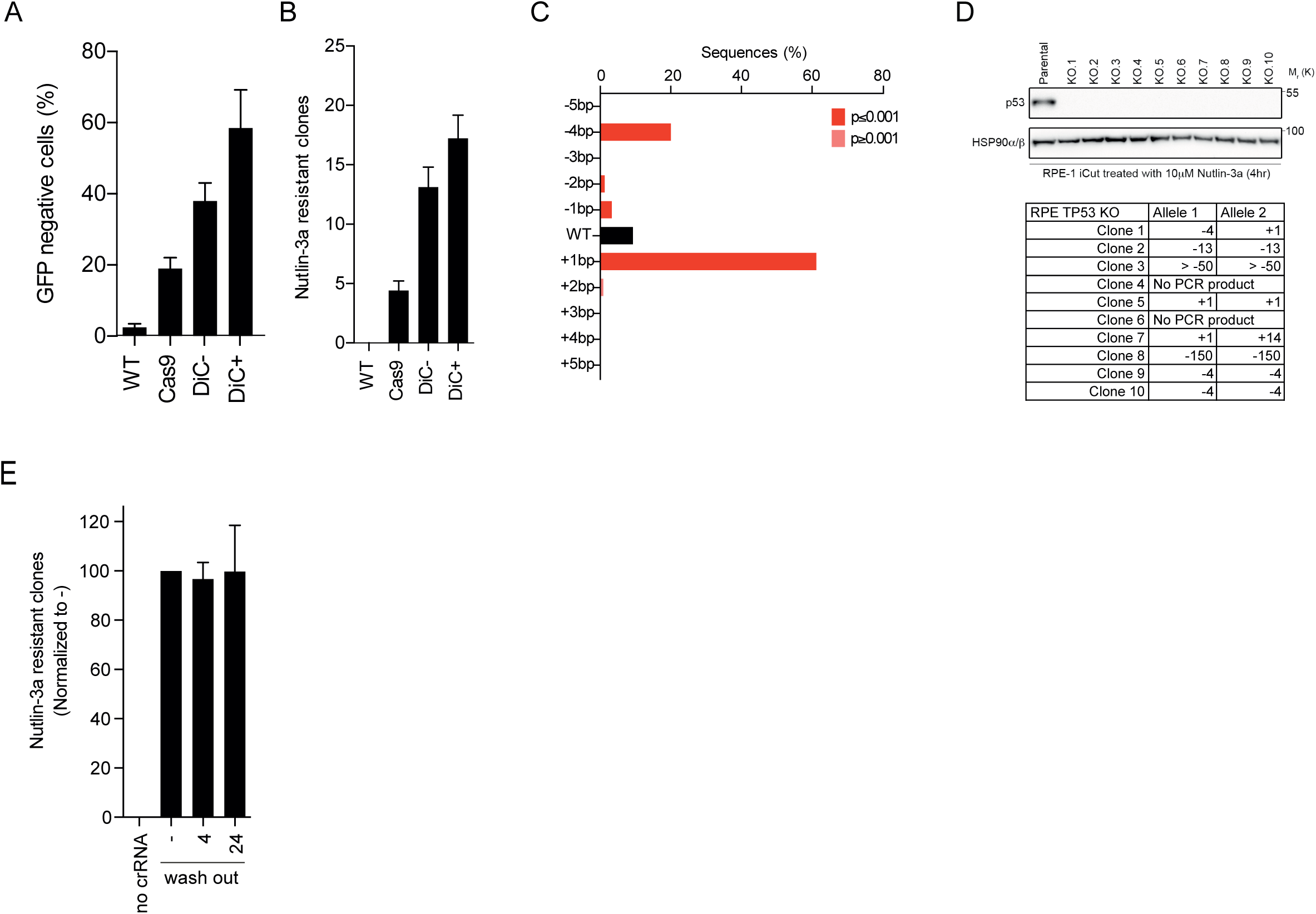
A) eGFP-assisted Cas9 activity assay with constitutive Cas9 expression of DiC cells with or without agonists. B) TP53/Nutlin-3a assisted Cas9 activity assay with constitutive Cas9 expression or DiC cells with or without agonists. C) Indel spectrum as readout with TIDE in iCut cells [27].D) Ten monoclonal TP53 knockout lines analyzed on protein level derived from iCut cells. E) Quantification of the number of Nutlin-3a selected clones to grow out in the RPE-1 DiC cell line. The average was determined from three independent experiments (error bars represent SEM).

**Supplemental Figure 3.**
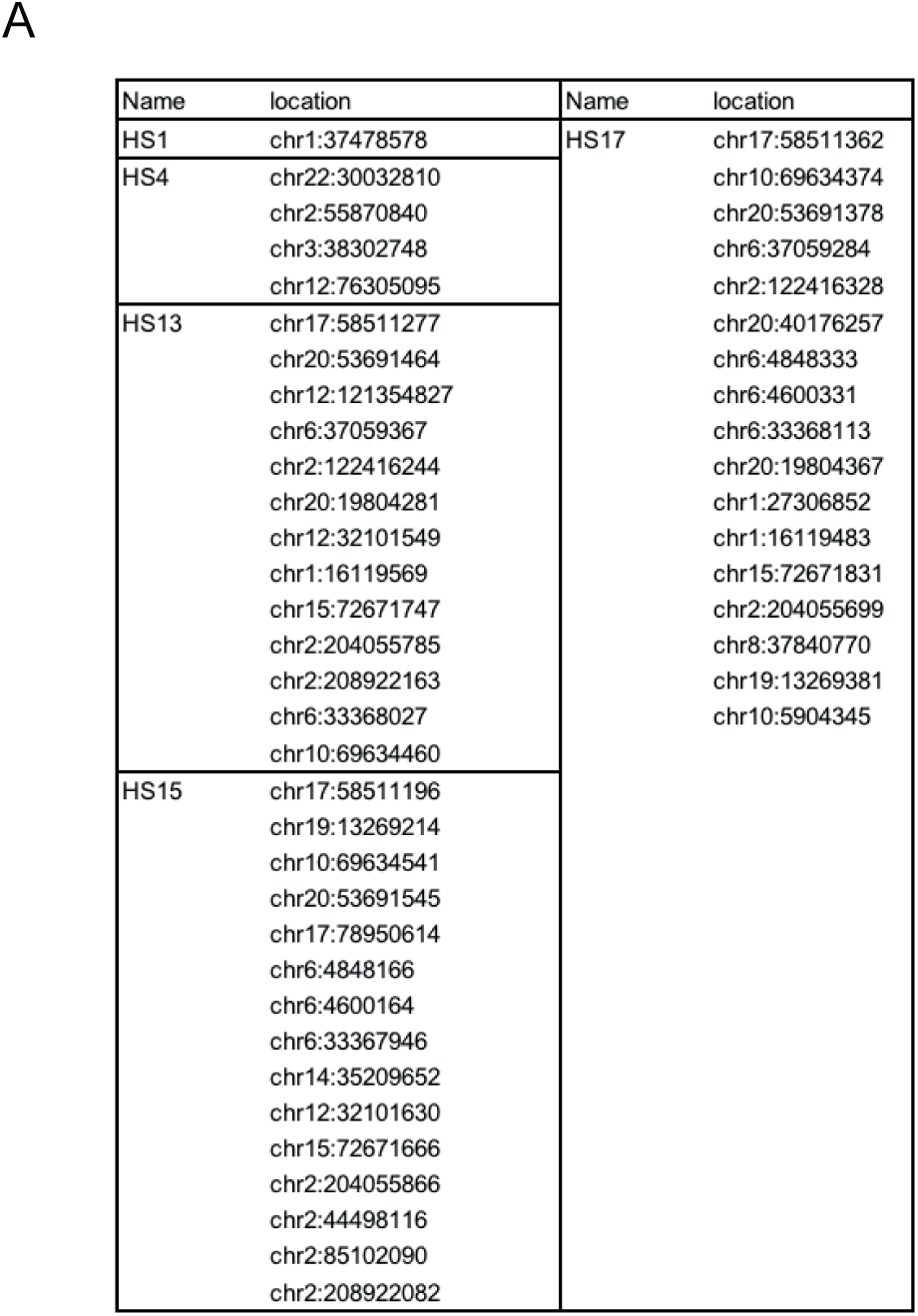
A) Target sites for gRNA used in Figs. 3-4.

**Supplemental Figure 4.**
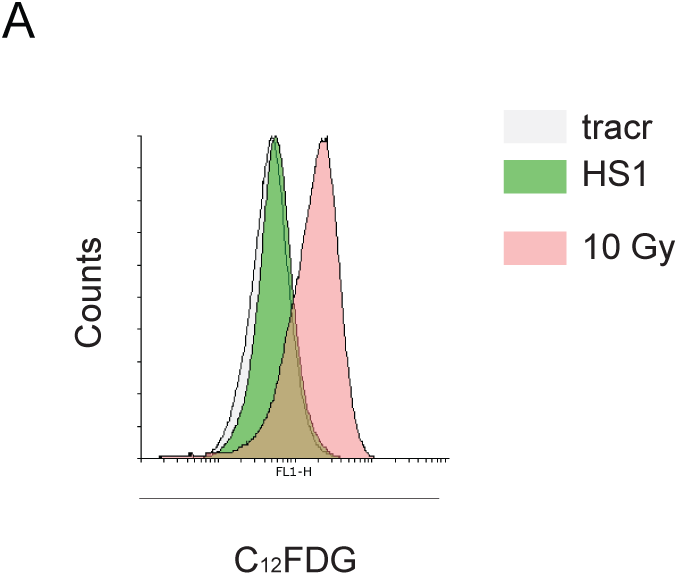
A) Cells were labeled with C_12_FDG 6 days after treatment or transfection and analyzed by Flow Cytometry. Cells were irradiated with 10 Gy (Pink), transfected with the tracr (Grey) or HS1 (Green).

